# Reanalysis of Orbitrap Astral DIA data demonstrates the capabilities of MS/MS-free proteomics to reveal new biological insights in disease-related samples

**DOI:** 10.1101/2024.04.01.587550

**Authors:** Mark V. Ivanov, Anna S. Kopeykina, Mikhail V. Gorshkov

## Abstract

Data-independent acquisition (DIA) became a method of choice for quantitative proteomics. With the advent of the combination of Orbitrap FTMS and asymmetric track lossless analyzer Astral these DIA capabilities were further extended with the recent demonstration of quantitative proteomic sample analyses at the speed of up to hundreds of samples per day. In particular, the dataset containing brain samples related to the multiple system atrophy was acquired using 7 and 28 min chromatography gradients and the Orbitrap Astral mass spectrometer (Guzman et al., Nat. Biotech.2024). In this work, we reanalysed the Orbitrap Astral DIA data using MS1 spectra without applying to fragmentation information using the recently introduced DirectMS1 approach. The results were compared with previous study of the same sample cohort by traditional long gradient DDA analysis. While the quantitation efficiency of DirectMS1 was comparable, we found additional five proteins of biological significance relevant to the analyzed tissue samples. Among the findings, DirectMS1 was able to detect decreased caspase activity for Vimentin protein in the multiple system atrophy samples which was barely observed from MS/MS-based methods. Our study suggests that DirectMS1 can be an efficient MS1-only addition to the analysis of DIA data in quantitative proteomic studies.

## Introduction

A recent study showed possibilities of a new combination of Orbitrap FTMS and asymmetric track lossless analyzer (Astral) in data-independent acquisition (DIA) for high-throughput proteomics with the capability to analyze up to 200 samples per day.^1^ This capability was demonstrated using brain tissue samples related to the multiple system atrophy analyzed previously by standard long-gradient label-free quantitative proteomics based on data-dependent acquisition method (DDA) using Orbitrap HF-X.^2^

In our previous study, we compared quantitation performance of our recently introduced DirectMS1Quant method with DIA-NN-based proteomic data analysis using the benchmark dataset consisting of a mix of *E. coli* proteome and UPS proteins at predefined concentrations.^3^ The comparison was performed for our own experimental data obtained using 5 min LC gradient MS1-only acquisition on FAIMS-Orbitrap Lumos mass spectrometer and publicly available DIA data for the same samples acquired earlier employing 90 min LC gradient on Orbitrap Lumos without FAIMS interface.^4^ That comparison demonstrated that the ultrafast proteomic analysis based on DirectMS1 method provides quantitation performance to the long-gradient DIA and significantly outperforms the latter for low concentration differentially expressed proteins present in the sample at the few fmol level.

Another important point derived from the mentioned comparison was that the efficient quantitation does not really depend on the number of identified and quantified proteins for a number of reasons. First, a majority of these proteins do not pass the significance thresholds when it comes to finding the differentially expressed one’s due to their underrepresentation by the corresponding peptides in MS/MS spectra. The latter is especially true in case of DDA analyses, where 50% of proteins are typically identified by one or two peptides. Second, some methods use non standard FDR control which makes comparison of any numbers between methods dubious. The latter is especially true in case of DIA analyses. For example, the popular DIA search engine DIA-NN uses a two-species library FDR approach or single N-terminal amino acid replacement for decoy sequence generation.^5^ Both approaches are not well-studied and widely-accepted in target-decoy practices routinely used by DDA search engines. And finally, there are many situations which could lead to wrong estimation of LFQ values even for true peptide/protein identifications.^6^

The Orbitrap Astral provides parallelized acquisition of MS1 and MS/MS spectra at the MS/MS scanning rate of ∼200 Hz and the isolation window down to 2 Th in DIA implementation, that positions this instrument as a method of choice in quantitative proteomics.^7^ However, the MS/MS-based quantitation, typically employed in DIA, has limitations, especially in the ultrashort LC gradient implementations and may miss, or underscore the important biological insights of the samples under study due to extremely high MS/MS signal interferences. At the same time, because of the parallelization, the proteomic data acquired using this novel instrument contain quite a number of MS1 spectra even from the short LC gradient analyses compared with the earlier versions of Orbitraps. It makes these data a valuable source for reanalysis using MS1 spectra which give researchers additional capabilities for revealing biologically relevant information from the same data otherwise uncovered in MS/MS spectra. There are several studies which explore the usage of MS1-based quantitation instead of MS/MS-based in DIA approaches.^8–11^ However, they all are about using MS/MS spectra for identification and MS1 signals for quantitation. Here, we performed the MS1-only spectra reanalysis of the Orbitrap Astral DIA data obtained for brain samples related to the multiple system atrophy (MSA), which were acquired using 7 and 28 min LC gradients.^1^ For the validation of the results we compared them with the ones reported in the recent study of the same sample cohorts by traditional 3h-long LC DDA.

## Methods

### Datasets

Data for the post-mortem brain tissue samples from the prefrontal cortex of 43 MSA patients and 27 normal controls were reanalysed in this study. The datasets were consisted of three groups: 180 min LC gradient DDA data obtained using Orbitrap Q-Exactive HF-X^2^; and Orbitrap Astral’s narrow-window DIA (nDIA) data obtained using two LC gradients, 7 and 28 min^1^. 7 min data were acquired in three technical replicates, while 28 min and 180 min in one only.

### DirectMS1 reanalysis

Raw files were converted into mzML format using ThermoRawFileParser^12^ (v. 1.3.4). Peptide features in MS1 spectra were detected using biosaur2^13^ (v. 0.2.19) which is freely available at https://github.com/markmipt/biosaur2 under Apache 2.0 license. Biosaur2 utility automatically extracts MS1 spectra from DIA data which are, then, used for detecting all peptide-like 13C isotopic clusters. For protein identification, the detected features were analyzed using MS1 search engine ms1searchpy^14^ (v. 2.7.3), also freely available at https://github.com/markmipt/ms1searchpy under Apache 2.0 license. Parameters for the search were as follows: 5% FDR, minimum 1 scan for detected peptide isotopic cluster; minimum one visible 13C isotope; charge states from 1+ to 6+, no missed cleavage sites, carbamidomethylation of cysteine as a fixed modification and 8 ppm initial mass accuracy, and peptide length range of 7 to 30 amino acid residues. DeepLC^15^ (v. 1.1.2) software was employed for predicting retention time of peptides, the information submitted to the ms1searchpy engine. DirectMS1Quant algorithm^3^ (distributed along with ms1searchpy) with default settings was used for detection of differentially expressed proteins which are 5% quantitative FDR with Benjamini Hochberg correction, two standard deviations of background proteins for fold change threshold and intensity normalization by 1000 quantified peptides with maximal intensities. Jupyter notebook with Python code to reproduce MS1-based analysis, starting from downloading raw files and ending with output tables with differentially expressed proteins, is available at https://github.com/markmipt/astral_brain_reanalysis/repository. It can be used as is without any modification on the Linux-based operating systems. Importantly, the code can be easily adopted by the researchers with minimal Python experience as it provides clear and straightforward instructions within the DirectMS1 workflow. Searches were performed against the Swiss-Prot human database, containing 20241 protein sequence. Decoy protein sequences for controlling FDR were generated by the pseudo shuffle method^3^ built-in ms1searchpy.

## Results and discussion

Note that despite the absence of ground truth knowledge of differentially expressed proteins and their real fold changes, the real-world samples with the known biological essence are the preferable platform for evaluating different quantitative proteomic methods compared with the in-lab prepared benchmarks containing pre-defined mixes of different species. The main reason for this is a simplicity of benchmark datasets due to a lack of high natural variability of proteins in the real-world samples significantly exceeding both technical and sample preparation variabilities.

General overview of the available datasets and identification results are shown in Figure 1. Expectedly, DirectMS1 analysis provided 4 times less identified proteins compared to DIA (1550 and 2300 for 7 and 28 min DirectMS1 vs 6600 and 8300 for 7 and 28 min DIA, respectively). Note that currently achievable depth of proteome DirectMS1 analysis using 5 min LC gradient on the FAIMS-Orbitrap Lumos at 120 000 MS1 resolution settings and three FAIMS CV values is ca. 3000 protein identifications at 5% FDR for 200 ng of commercial HeLa standard. As for the number of “quantified” proteins, we found across the literature too large a variety of definitions, mostly depending on the method of proteome analysis employed. Commonly, it is defined as the number of identified proteins in at least N samples, with the coefficient of variation less than 20% and percentage of missing values less than some threshold. In the DirectMS1 method we report the number of proteins which were identified in at least 1 sample and have at least 3 peptides with at least 50% non-missed intensities. At the same time, we believe that the popular reporting purely the number of “identified and quantified” proteins, as well as the coefficients of variation provide no warranty for better quantitation results in comparable methods, or tell much about the perspectives of obtaining biologically important information about regulated proteins and/or cellular processes.^3,6^

**Figure 1.**
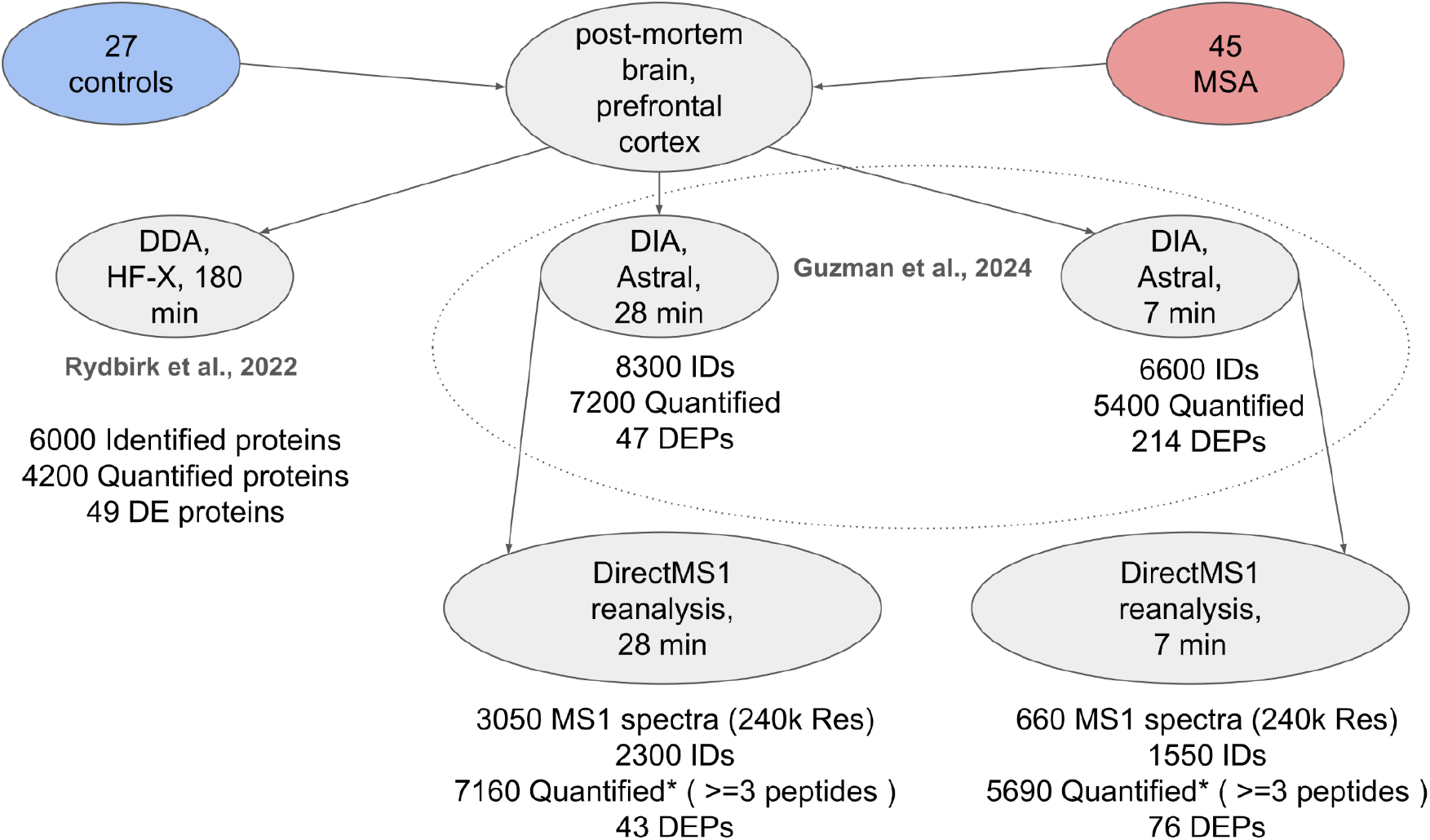
Overview of datasets and results obtained using different proteomic analysis methods.The number of differentially expressed proteins for DIA results and all numbers for DirectMS1 reanalysis were obtained in this study. The rest of the values were extracted from the original manuscripts. *Note, that the number of quantified proteins is a union for all runs without additional FDR control. DirectMS1Quant is able to control quantitative FDR despite a high level of identification FDR, see Supplementary Figure S2 in previous study^3^.

### Differentially expressed proteins

DirectMS1Quant reported 76 and 43 differentially expressed proteins (DEPs) for 7 and 28 min data, respectively (Figure 2a). These numbers were obtained using a fold change threshold estimated by DirectMS1Quant as two standard deviations of the background proteins in the sample, and the p-value threshold which was 0.05 with Benjamini–Hochberg correction. While the list of DEPs for DIA analysis were not provided in the referenced manuscript, one can recognize approx. 35 and 20 DEPs in Fig.6d for 7 and 28 min data, respectively. For comparison with MS1-only results, we applied our own fold change and p-value calculations using LFQ values presented in the Supplemental tables of the above manuscript. By applying 0.28 and 0.44 log_2_(FC) thresholds for 7 and 28 min data, these calculations resulted in 214 and 47 DEPs for 7 and 28 min data, respectively. Note that both p-values and fold change thresholds cannot be directly comparable between the methods and the higher number of DEPs itself does not mean a better quantitation.

**Figure 2.**
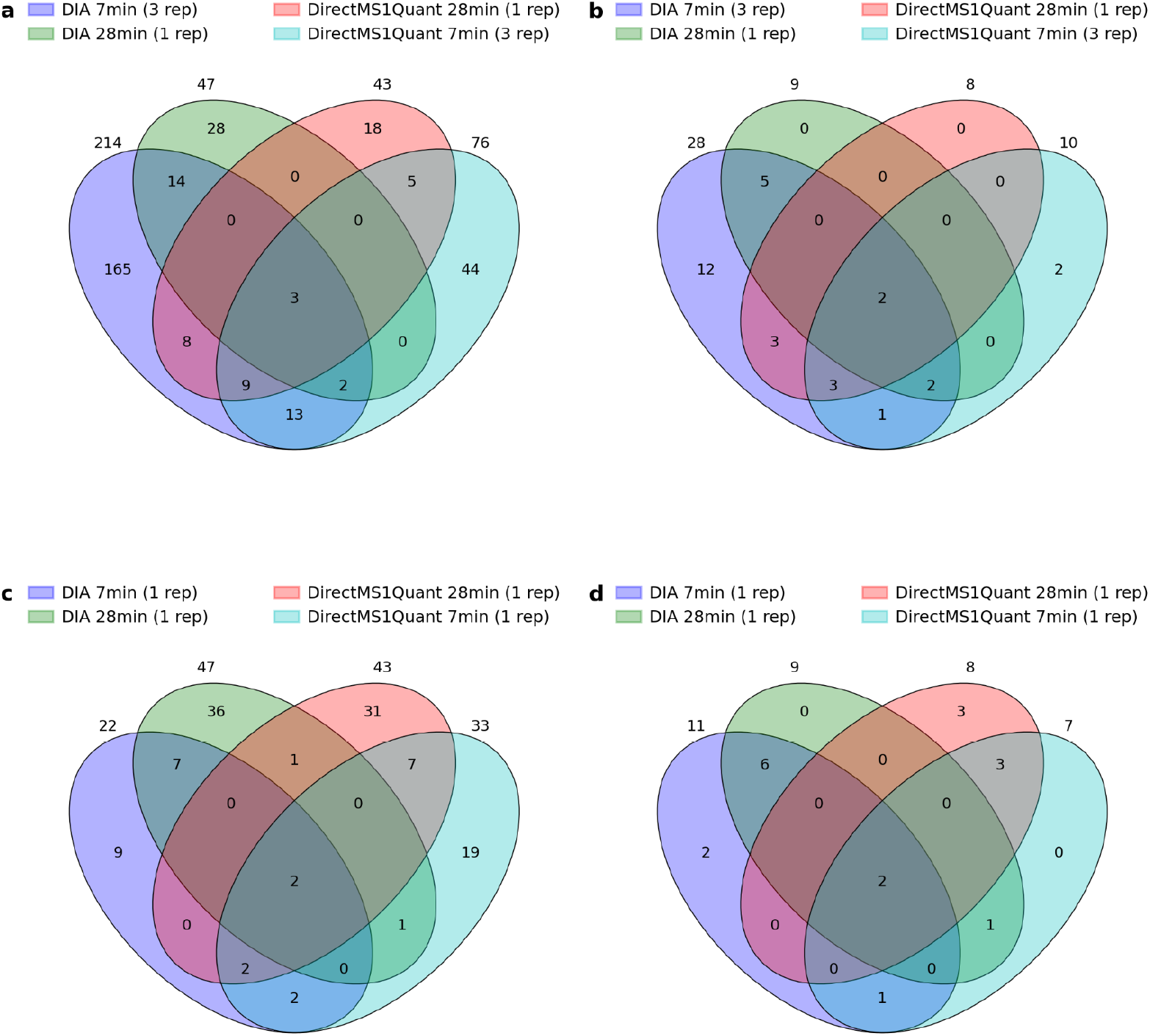
Venn diagrams for reported differentially expressed proteins for: (a,b) - 7 min DirectMS1 (in triple replicates), 28 min DirectMS1, 7 min DIA (in triple replicates) and 28 min DIA data; and (c,d) - 7 min data were used in single technical replicate. (a,c) - all reported DE proteins; (b, d) - reported DE proteins confirmed by work based on 180 min LC DDA data.

Next, we performed validating the results provided by DirectMS1Quant and DIA. First, we looked at the intersection between our results and DEPs reported earlier for the same samples using long-gradient deep proteome DDA analysis. As shown in Figure 2b, 10, 8, 29, and 9 DEPs found in MS1 7 min, MS1 28 min, DIA 7 min, and DIA 28 min data, respectively, were confirmed by DDA. Surprisingly enough, DIA 7 min analysis outperforms all other methods, in spite of that 28 min DIA resulted in more identified and quantified proteins. This observation further supports our viewpoint on the overstated importance of the number of IDs compared with the number of DEPs when it comes to expression proteomics applications. The number of replicates is more important for quantitative proteomics, which further stresses the importance of developing high-throughput acquisition methods. Indeed, as shown in this and referenced studies the difference between 7 and 28 min results within one method, DIA or DirectMS1, can be easily explained by using triple replicates when acquiring 7 min data. Indeed, the number of differentially expressed proteins from a single 7 min replicate are shown in Fig. 2c which is way below the results from a single 28 min replicate for both methods. Further comparison of triple 7 min replicate with single 28 min replicate shows that spending more instrument time for technical replicates is more useful for quantitation than for longer gradients. In the context of the above discussion, it would be quite logically to ask, how many of DEPs reported by short gradient DIA and DirectMS1 methods and then confirmed by long gradient DDA analysis of the same samples? This is shown in Figs. 2b and 2d. Again, shorter gradient allowing performing more technical replicates results in more confirmed DEPs in spite of smaller number of IDs. Importantly to note here, that DirectMS1 applied to 28 min data produced a similar number of DEPs as 28 min DIA as shown in Figs.2c and 2d. This is not the case for 7 min data, Figs.2a and 2b, because the number of extracted MS1-only spectra from these data were only ca. 660. Speaking of DirectMS1 performance when data are acquired in MS1-only mode this number of MS1 spectra would correspond to 1-2 min gradient analysis on state-of-the art Orbitrap MS at 120k resolution (see, more detailed DirectMS1 method description elsewhere.^16^ Yet, even in this case, the numbers of DEPs confirmed by DDA analysis were comparable. From this viewpoint, it would be fairer to compare single 28 min replicate DirectMS1, which corresponds to 7 min MS1-only acquisition in terms of the number of obtained MS1 spectra, with single 7 min replicate DIA. As shown in Figs. 2c, DirectMS1 outperforms in the number of reported DEPs. Two proteins, P02675 (Fibrinogen beta chain) and P02679 (Fibrinogen gamma chain), were confirmed by all 5 quantitation methods (7 min and 28 min DirectMS1, 7 and 28 min DIA, and 180 min DDA) (see Supplementary Table 1). It was not surprising, as the previous DDA-based study has shown that fibrinogen accumulation is one of the key processes associated with MSA. In addition, one more fibrinogen protein, P02671 (Fibrinogen alpha chain) was reported as differential expressed by four quantitation methods except the 28 min DIA. However, the latter provided fold change estimation close to the other methods (log2(FC)=0.37; uncorrected p-value of 0.014), and this protein marginally missed both fold change and p-value thresholds.

To further elaborate on this issue, note that 120k MS1 spectra resolution provides a better number of identified proteins for the DirectMS1 method compared with the 240k resolution used in the referenced Astral’s work. Moreover, we expect that the difference in quantitation results in terms of reported DEPs may be even more dramatic. Indeed, among 660 MS1 extracted from the Orbitrap Astral 7 min data, only approximately 550 contained peptide signals (Fig. 3a). The distribution of the number of MS1 scans per matched peptide is shown in Fig. 3b. Most of the detected peptide isotopic clusters are presented only in one, two, or three scans. Despite that, completeness for peptides belonging to reported DEPs across the samples is still good (Fig. 3c). The distribution of the number of MS1 scans for those peptides is shown in Fig. 3d. We can see practically no peptides with 1 or 2 MS1 scans per profile for the reported DEPs. This means that one needs more than 3 MS1 scans per profile to extract peptide intensity apex accurately enough for a protein to pass statistical thresholds in the further quantitation analysis. Thus, one should expect that 7 min data acquisition with 120k as we usually do for DirectMS1 method would be closer to the results obtained here for the 28 min LC with 240k due to higher coverage of peptide elution profiles with MS1 spectra. The above considerations allow expecting, even for the Orbitrap MS of previous generations, close or even better quantitation results when using DirectMS1 method in MS1-only acquisition mode with triple 7 min replicates of 120k resolution per MS1 spectra compared with DIA analysis of the same duration .

**Figure 3.**
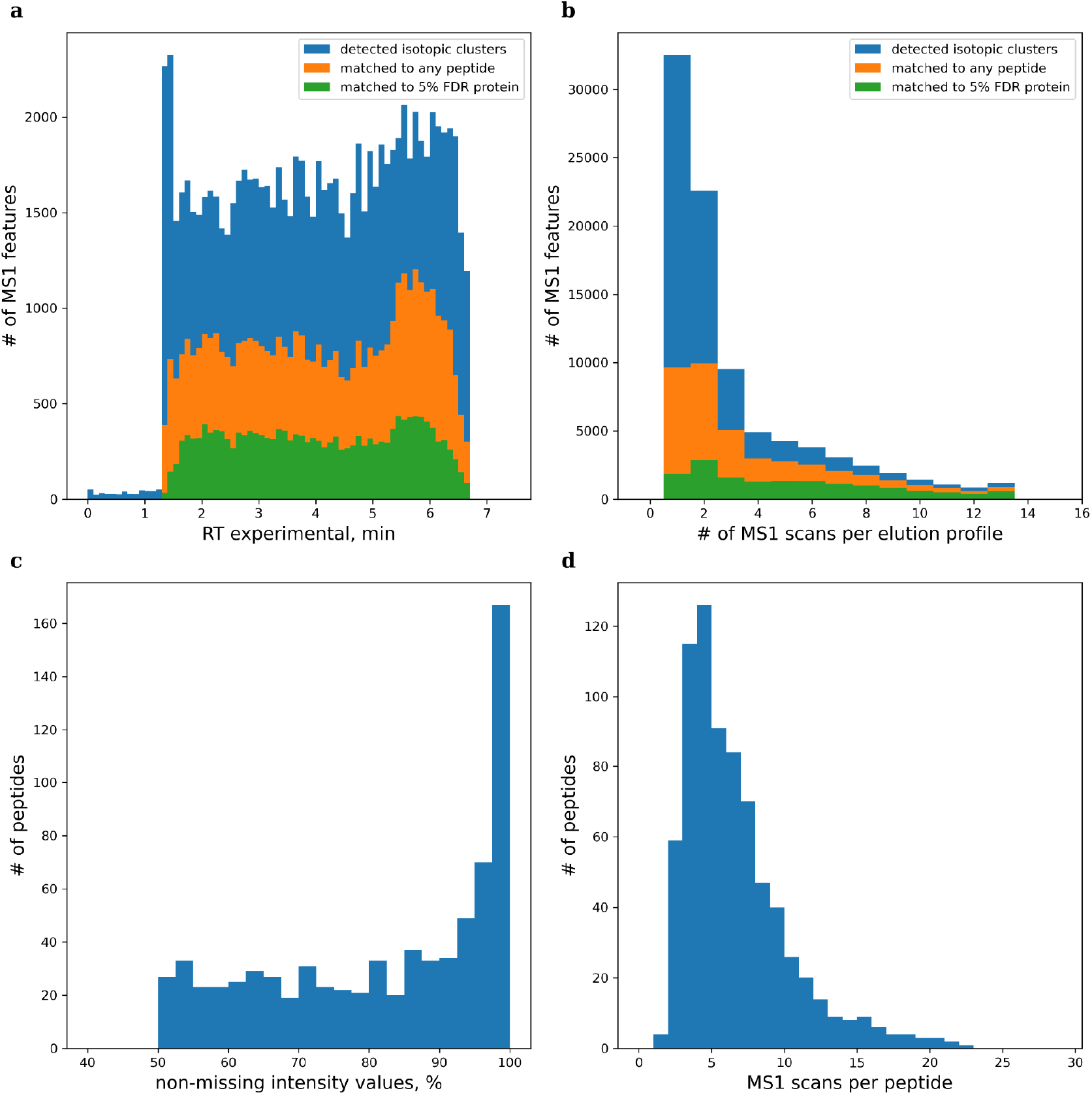
**(a, b)** Distribution of detected peptide isotopic clusters by (a) experimental retention time and (b) number of MS1 scans pep peptide elution profile. Blue color shows all detected MS1 features; orange color shows those which are matched to any peptide in the database by ms1searchpy; and green color shows those which are matched to peptides belonging to the list of proteins identified at 5% FDR. The data is shown for a randomly chosen file from the data set. **(c, d)** Distribution of quantified peptides in DirectMS1 analysis of 7 min LC data by (c) non-missing intensity values across all analyzed files; and (d) average number of MS1 scans per peptide elution profile for peptides belonging to reported differentially expressed proteins.

### Analysis of novel DirectMS1-only proteins

We found 5 proteins which were reported as DEPs only by DirectMS1: P02792 (Ferritin light chain), P06576 (ATP synthase subunit beta, mitochondrial), P08670 (Vimentin), Q14CZ8 (Hepatic and glial cell adhesion molecule) and Q6YN16 (Hydroxysteroid dehydrogenase-like protein 2) (Table 1). Note that these proteins were identified and quantified in DIA analysis although their estimated fold changes were twice less compared to the ones reported by DirectMS1. Interestingly, the DIA analysis returned twice more quantified peptides for those proteins compared with DirectMS1, and probably the variations of their intensities in the MS/MS-based analysis precluded them from passing statistical significance threshold by p-value. In the following, we performed a literature search to prove that these findings make sense from the biological viewpoint rather than just false reports due to our method’s flaws.

**Table 1.** List of proteins reported by DirectMS1 analysis as differentially expressed in both 7 and 28 min datasets and missed in MS/MS-based analyses. p-values are provided in uncorrected form.

| dbname | description | 28 min data |  |  |  | 7 min data |  |
| --- | --- | --- | --- | --- | --- | --- | --- |
|  |  | # of quantified peptides |  | log<sub>2</sub>(FC)<br>(MSA/Control) /<br>p-value |  | log<sub>2</sub>(FC)<br>(MSA/Control) /<br>p-value |  |
|  |  | DirectMS1 | DIA | DirectMS1 | DIA | DirectMS1 | DIA |
| P02792 | Ferritin light chain | 6 | 13 | -1.38 /<br>3.9E-5 | -0.8 /<br>1.8E-1 | -1.01 /<br>1.9E-4 | -0.54 /<br>3.3E-2 |
| P06576 | ATP synthase subunit beta, mitochondrial | 21 | 41 | -0.72 /<br>2.3E-3 | -0.12 /<br>2.5E-1 | -0.28 /<br>3.9E-20 | -0.15 /<br>7.2E-3 |
| P08670 | Vimentin | 31 | 70 | 0.6 /<br>6.1E-4 | 0.22 /<br>5.5E-2 | 0.37 /<br>2.7E-10 | 0.23 /<br>2.1E-4 |
| Q14CZ8 | Hepatic and glial cell adhesion molecule | 10 | 27 | -0.64 /<br>2.3E-4 | -0.25 /<br>6.4E-3 | -0.34 /<br>1.9E-4 | -0.16 /<br>8.0E-4 |
| Q6YN16 | Hydroxysteroid dehydrogenase-like protein 2 | 17 | 40 | -0.47 /<br>1.2E-3 | -0.12 /<br>3.4E-2 | -0.38 /<br>3.2E-4 | -0.1 /<br>5.0E-6 |

### P02792 protein

Anomalous metal distribution is characteristic of both neurodegenerative diseases in general and alpha-synucleinopathies, such as multiple system atrophy in particular.^17–19^ Hence, examining the pathogenesis of this disease specifically focuses on the increase in iron concentration in subcortical brain structures in combination with the imbalance of iron metabolism regulator proteins.^20^ Our analysis of samples from the prefrontal cortex revealed a decrease in the level of the protein P02792 (light chain of ferritin) relative to controls, as was previously demonstrated immunohistochemically in human brain samples.^21^ However, in previous studies, statistically significant differences in ferritin levels in human brain tissues^22^ and mouse model lines^23^ were not observed, although there was a tendency towards a decrease in protein levels. Given that iron levels in this brain area are stable^21,22^, one can indirectly infer an increase in free bioavailable iron in the prefrontal cortex of MSA patients. This can contribute to increased oxidative stress due to the production of reactive oxygen species (ROS). ROS, in turn, can damage DNA and mtDNA, carbonylate proteins, inducing lipid peroxidation with cell membrane damage.^24^ Additionally, it has been shown that *in vitro* aggregation of α-synuclein is triggered by increased concentrations of trivalent iron.^25^ These processes collectively form a ‘vicious cycle’ that contributes to the death of neural cells and the progression of MSA. A broader understanding of iron homeostasis in various brain structures in such patients may shed light on the mechanisms underlying the development of this disease and establish accurate causal relationships in molecular cascades.

### P08670 and Q14CZ8 proteins

Changes in the expression of other proteins, such as P08670 (Vimentin) and Q14CZ8 (hepatic and glial cell adhesion molecule), can be linked to neuroinflammation caused by cytoplasmic cellular inclusions, glial activation, and neural cell apoptosis.^26–28^ It has been shown that the astrogliosis developing in MSA affects both cortical and subcortical structures.^29–31^ Such activation of the immune system increases the number of reactive astrocytes, which widely express mesenchymal markers.^32^ This assertion is supported by other differentially expressed proteins reported in our study (Supplementary Table 1), where an increase in several mesenchymal markers (VIM+, FIBG+, FIBB+) and a decrease in the expression of epithelial markers (GJB6-, MATR3-) can be observed. Moreover, the increase in Vimentin expression in MSA was also previously demonstrated in transcriptomic analysis.^33^ Limited literature exists on the protein Q14CZ8, which is primarily mentioned in the context of various types of cancer, rather than the neurodegenerative diseases. For example, it has been shown to inhibit proliferation, migration, and invasion of cancer cells.^34,35^ Extrapolating these findings to neural tissue, it can be said that the number of mesenchymal markers increases due to gliosis activation in MSA.

### P06576 protein

Based on the literature, one of the components of MSA pathogenesis is dysfunction of the electron transport chain, resulting in increased oxidative stress due to reduced COQ2 expression, leading to the corresponding reductions in CoQ and ATP levels in the motor cortex, cerebellum, and pons.^36–38^ According to our findings, the decrease in ATP levels in the prefrontal cortex may be associated with ATP synthase dysfunction due to decreased expression of the protein P06576 (ATP synthase subunit beta). Changes in the expression of ATP synthase subunits can also disrupt cellular energy metabolism and lead to mitochondrial dysfunction, as seen in other neurodegenerative diseases.^39^

### Q6YN16 protein

The protein Q6YN16 (hydroxysteroid dehydrogenase-like protein 2 - HSDL2) is involved in fatty acid metabolism and is well associated with the progression of various types of cancer.^40,41^ For example, it has been shown that in various types of gliomas, overexpression of this protein induces proliferation and increases cell survival, while gene knockdown leads to apoptosis and cell cycle arrest.^42^ In contrast to the previous study, a recent report demonstrated that knockdown of HSDL2 led to inhibition of ferroptosis in cholangiocarcinoma cells, thereby promoting cancer progression.^43^ This was achieved through the regulatory factor p53, which is also active in neurodegeneration.^44^ In this regard, it can be hypothesized that the reduction of HSDL2 expression in MSA acts as neuroprotection for brain cells prone to ferroptosis.^45^

Interestingly, a number of proteins typically associated with MSA (α-Synuclein, Ubiquitin, α- and β-Tubulins, αB-Crystallin, Tau, MAPs proteins)^46^ were not found to be differentially expressed proteins in both our and original DDA/DIA studies, which can be explained by the region from which the samples were taken. However, based on our results, we can infer molecular changes affecting the prefrontal cortex, including regulation of iron metabolism, mitochondrial dysfunction with increased oxidative stress, induction of ferroptosis, and neuroinflammation. Based on the existing literature, one can conclude that complex multilevel cascades exist and their functioning levels vary among different patients with different forms of MSA. Therefore, a more detailed investigation is necessary, including experimental groups categorized by both clinical criteria and brain regions. A more comprehensive review of the issue can be found elsewhere.^47–51^

### More on the Vimentin protein in view of caspase activity

During manual inspection of DEPs reported by DirectMS1 method, we noticed that most valuable information showing differential expression of Vimentin protein comes from peptides from the middle (104-236 amino acid positions) and last part (294-410) of the protein sequence (Supplementary Table 2). Literature search^52^ showed that multiple caspases cleave Vimentin at Asp sites at the positions 85 and 259, which can explain our observations. Also, according to the MEROPS^53^ database with a reference to Julien et al. study^54^, Vimentin can be also cleaved at Asp at position 90. We expanded our database with three semi-tryptic peptide pairs, LLQDSVD + FSLADAINTEFK; LLQDSVDFSLAD + AINTEFK; LHEEEIQELQAQIQEQHVQIDVD + VSK for the cleavage at positions 85, 90 and 259, respectively, and repeated DirectMS1 analysis. Semi-tryptic peptide intensities were extracted and normalized by the sum of all fully tryptic peptides belonging to Vimentin protein. Peptide FSLADAINTEFK (Asp85 cleavage) clearly shows a decrease in normalized intensity for MSA patients samples which is in contrast with general increase of Vimentin protein for the MSA group (Figure 4a). This counter move of the concentration of the protein and one of its peptides is one of the important indications of ferment activities. Its paired peptide AINTEFK was detected only in 28 min DIA data and shows the same fold change direction but with no significant difference (Fig. 4b). Also, we were able to detect difference in expression of peptide AINTEFK (Fig. 4c), but the statistical significance and changes are much smaller compared to the cleavage at Asp85 observed for FSLADAINTEFK. The rest of semi-tryptic peptides were detected in some runs, but we assume that they are mostly false matches. All mentioned results have a low chance of coincidence despite the fact that DirectMS1 method do not control peptide-level FDR. To further validate our findings and prove that these are not just MS1 features belonging to any protein whose level is decreased in case of MSA samples, we made a classic DDA closed search using the MSFragger^55^ search engine. The search parameters were standard except precursor and fragment mass tolerances were set at 10 ppm and 0.02 Da, respectively. Also, the isotopic error parameter was turned off to decrease search space and to obtain more conservative results. Peptide FSLADAINTEFK was identified only in 4 of 75 DDA files and all MS/MS spectra had different retention times. Only one identified MS/MS spectra had RT (119.54 min) corresponding exactly to the MS1 peptide feature reported by biosaur (starting from 119.48 min, ending 119.69 min, intensity apex at 119.61 min) and identified as FSLADAINTEFK peptide in DirectMS1 workflow. Also FSLADAINTEFK was matched by 15 fragment ions of 22 theoretical ones which is considered as an extremely good coverage (Fig. 4d). More interesting is that 3 hour DDA analysis was not able to detect this peptide in 74 of 75 experimental runs. Peptide AINTEFK was identified in 31 DDA runs with 8 matched ions of 12 theoretical fragments. Peptide LLQDSVD was not identified in DDA analysis at all.

**Figure 4.**
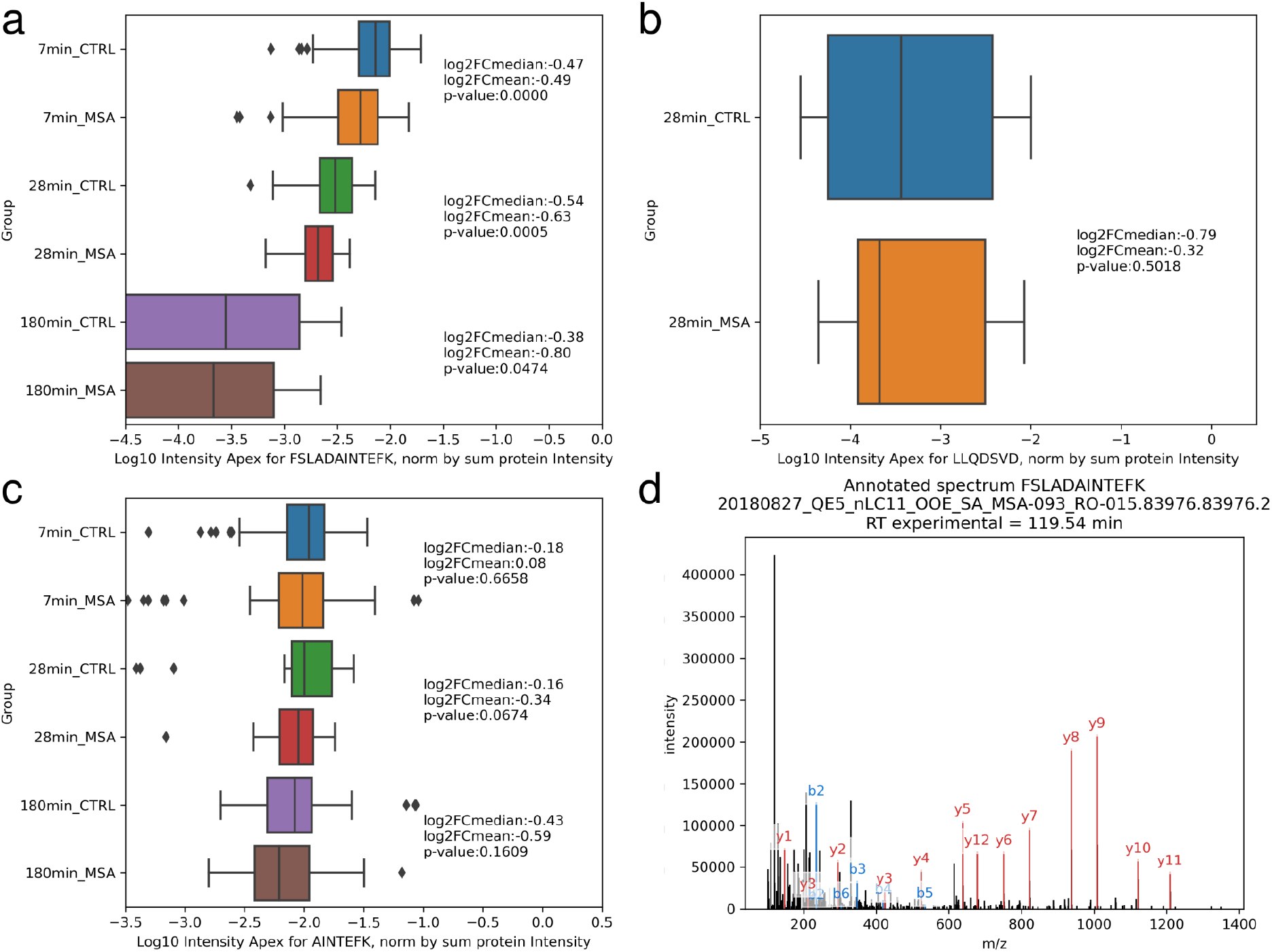
(a-c) Boxplots with intensities of different semi-tryptic peptides of Vimentin protein for 3 data sets: 7 min and 28 min DIA Astral data and 180 min DDA HF-X data. Fold changes are calculated as an MSA group divided by the Control group. Peptide intensities were normalized by the sum of all peptide intensities belonging to Vimentin protein. p-values were calculated using independent t-test with no correction. (d) Matched MS/MS spectrum of FSLADAINTEFK peptide.

It has long been shown that cleavage of Vimentin at the sites described above signals on the activity of the mitochondrial-dependent apoptosis pathway through the activation of initiator and effector caspases and the assembly of the apoptosome.^56^ Vimentin is one of the substrates for effector caspases 3, 6, and 7, and its cleavage products serve as a pro-apoptotic signal leading to cell death.^52^ Based on our data, it can be concluded that the activity of effector caspases 3, 7, and 6 towards Vimentin protein is reduced in MSA. The reasons for such functional changes are speculative. On the one hand, many publications report an increased activity of both initiator and effector caspases, as well as proteins involved in apoptosome assembly.^57–60^ On the other hand, the activity of these proteins has been associated with intracellular inclusions from subcortical brain structures^59^ or data obtained from cellular models^57,58^. Of course, there is no doubt about the progressive development of neuroinflammation in subcortical brain structures in MSA. Perhaps the decrease in the mitochondrial-mediated apoptosis pathway in cells of the prefrontal cortex is associated with the activity of other forms of cell death, such as ferroptosis or autophagy.^45,61^ Alternatively, effector caspases may change their substrate specificity and/or activity to protect cells from death in as yet unaffected parts of the brain. The question also remains open: are our data related to the specificity of cell material collection and the region from which it was taken, or do they represent incompletely understood biological processes?

## Conclusions

Over the last few decades, mass-spectrometry based proteomics is focusing on optimization of fragmentation methods for protein identification and quantitation and there is no exception for the recently introduced state-of-art Orbitrap Astral. This work validates our earlier attempts to demonstrate a potential for MS1-only spectra processing methods for proteomics which are rarely considered as standalone solutions for protein expression analysis. Indeed, this work shows that 3000 of MS1 spectra is enough to extract quantitative information about the proteins similar to the one attainable from 400 000 tandem mass spectra, which can be acquired by the novel Orbitrap Astral MS operated at the maximum acquisition rate of 200 Hz in a half an hour analysis. Note that ultrafast proteome-wide analysis in a minute time scale is the primary avenue for DirectMS1 method that make it well-suited for biomedical and clinical applications, in which quantitative capabilities of a method and, often, its simplicity are the important issues for consideration. However, we show in this work that the MS1-only spectra reanalysis of previously acquired MS/MS-based proteomic data, either DIA, or DDA, may bring new biological insights on the samples under study. Specifically, the method identified additional five differentially expressed proteins which can be associated with MSA disease, while these proteins were missed in MS/MS-based DIA quantitation. Note also on the principal difference between DirectMS1 and MS1 spectra quantitation currently employed in proteomics. The latter relies on the intensities of peptide ions in MS1 spectra which were found to match the underlying MS/MS spectra. DirectMS1 takes into account all peptide-like features detectable in MS1 spectra and uses them all for protein identification and quantitation that adds quantitation power to the method.

## Acknowledgments

Study was performed with financial support from the Russian Science Foundation, grant no. 23-45-00012 to M.V.G.

## Authors contributions

M.V.I. proposed the concept of the study and processed all mass-spectrometry data. M.V.I. and A.S.K. critically evaluated the results. A.S.K. done biomedical research of detected proteins. M.V.I. and M.V.G. wrote the first draft of the paper. All authors read, edited and approved the final version of the paper.

## References

(1) Guzman, U. H.; Martinez-Val, A.; Ye, Z.; Damoc, E.; Arrey, T. N.; Pashkova, A.; Renuse, S.; Denisov, E.; Petzoldt, J.; Peterson, A. C.; Harking, F.; Østergaard, O.; Rydbirk, R.; Aznar, S.; Stewart, H.; Xuan, Y.; Hermanson, D.; Horning, S.; Hock, C.; Makarov, A.; Zabrouskov, V.; Olsen, J. V. Ultra-Fast Label-Free Quantification and Comprehensive Proteome Coverage with Narrow-Window Data-Independent Acquisition. Nat. Biotechnol. 2024, 1–12. 10.1038/s41587-023-02099-7.

(2) Rydbirk, R.; Østergaard, O.; Folke, J.; Hempel, C.; DellaValle, B.; Andresen, T. L.; Løkkegaard, A.; Hejl, A.-M.; Bode, M.; Blaabjerg, M.; Møller, M.; Danielsen, E. H.; Salvesen, L.; Starhof, C. C.; Bech, S.; Winge, K.; Rungby, J.; Pakkenberg, B.; Brudek, T.; Olsen, J. V.; Aznar, S. Brain Proteome Profiling Implicates the Complement and Coagulation Cascade in Multiple System Atrophy Brain Pathology. Cell. Mol. Life Sci. 2022, 79 (6), 336. 10.1007/s00018-022-04378-z.

(3) Ivanov, M. V.; Bubis, J. A.; Gorshkov, V.; Tarasova, I. A.; Levitsky, L. I.; Solovyeva, E. M.; Lipatova, A. V.; Kjeldsen, F.; Gorshkov, M. V. DirectMS1Quant: Ultrafast Quantitative Proteomics with MS/MS-Free Mass Spectrometry. Anal. Chem. 2022, 94 (38), 13068–13075. 10.1021/acs.analchem.2c02255.

(4) Gotti, C.; Roux-Dalvai, F.; Joly-Beauparlant, C.; Mangnier, L.; Leclercq, M.; Droit, A. Extensive and Accurate Benchmarking of DIA Acquisition Methods and Software Tools Using a Complex Proteomic Standard. J. Proteome Res. 2021, 20 (10), 4801–4814. 10.1021/acs.jproteome.1c00490.

(5) Demichev, V.; Messner, C. B.; Vernardis, S. I.; Lilley, K. S.; Ralser, M. DIA-NN: Neural Networks and Interference Correction Enable Deep Proteome Coverage in High Throughput. Nat. Methods 2020, 17 (1), 41–44. 10.1038/s41592-019-0638-x.

(6) Ivanov, M. V.; Garibova, L. A.; Postoenko, V. I.; Levitsky, L. I.; Gorshkov, M. V. On the Excessive Use of Coefficient of Variation as a Metric of Quantitation Quality in Proteomics. PROTEOMICS 2024, 24 (1–2), 2300090. 10.1002/pmic.202300090.

(7) Stewart, H. I.; Grinfeld, D.; Giannakopulos, A.; Petzoldt, J.; Shanley, T.; Garland, M.; Denisov, E.; Peterson, A. C.; Damoc, E.; Zeller, M.; Arrey, T. N.; Pashkova, A.; Renuse, S.; Hakimi, A.; Kühn, A.; Biel, M.; Kreutzmann, A.; Hagedorn, B.; Colonius, I.; Schütz, A.; Stefes, A.; Dwivedi, A.; Mourad, D.; Hoek, M.; Reitemeier, B.; Cochems, P.; Kholomeev, A.; Ostermann, R.; Quiring, G.; Ochmann, M.; Möhring, S.; Wagner, A.; Petker, A.; Kanngiesser, S.; Wiedemeyer, M.; Balschun, W.; Hermanson, D.; Zabrouskov, V.; Makarov, A. A.; Hock, C. Parallelized Acquisition of Orbitrap and Astral Analyzers Enables High-Throughput Quantitative Analysis. Anal. Chem. 2023, 95 (42), 15656–15664. 10.1021/acs.analchem.3c02856.

(8) Huang, T.; Bruderer, R.; Muntel, J.; Xuan, Y.; Vitek, O.; Reiter, L. Combining Precursor and Fragment Information for Improved Detection of Differential Abundance in Data Independent Acquisition*. Mol. Cell. Proteomics 2020, 19 (2), 421–430. 10.1074/mcp.RA119.001705.

(9) Schilling, B.; Rardin, M. J.; MacLean, B. X.; Zawadzka, A. M.; Frewen, B. E.; Cusack, M. P.; Sorensen, D. J.; Bereman, M. S.; Jing, E.; Wu, C. C.; Verdin, E.; Kahn, C. R.; MacCoss, M. J.; Gibson, B. W. Platform-Independent and Label-Free Quantitation of Proteomic Data Using MS1 Extracted Ion Chromatograms in Skyline: APPLICATION TO PROTEIN ACETYLATION AND PHOSPHORYLATION*. Mol. Cell. Proteomics 2012, 11 (5), 202–214. 10.1074/mcp.M112.017707.

(10) Rardin, M. J.; Schilling, B.; Cheng, L.-Y.; MacLean, B. X.; Sorensen, D. J.; Sahu, A. K.; MacCoss, M. J.; Vitek, O.; Gibson, B. W. MS1 Peptide Ion Intensity Chromatograms in MS2 (SWATH) Data Independent Acquisitions. Improving Post Acquisition Analysis of Proteomic Experiments*[S]. Mol. Cell. Proteomics 2015, 14 (9), 2405–2419. 10.1074/mcp.O115.048181.

(11) Egertson, J. D.; Kuehn, A.; Merrihew, G. E.; Bateman, N. W.; MacLean, B. X.; Ting, Y. S.; Canterbury, J. D.; Marsh, D. M.; Kellmann, M.; Zabrouskov, V.; Wu, C. C.; MacCoss, M. J. Multiplexed MS/MS for Improved Data-Independent Acquisition. Nat. Methods 2013, 10 (8), 744–746. 10.1038/nmeth.2528.

(12) Hulstaert, N.; Shofstahl, J.; Sachsenberg, T.; Walzer, M.; Barsnes, H.; Martens, L.; Perez-Riverol, Y. ThermoRawFileParser: Modular, Scalable, and Cross-Platform RAW File Conversion. J. Proteome Res. 2020, 19 (1), 537–542. 10.1021/acs.jproteome.9b00328.

(13) Abdrakhimov, D. A.; Bubis, J. A.; Gorshkov, V.; Kjeldsen, F.; Gorshkov, M. V.; Ivanov, M. V. Biosaur: An Open-Source Python Software for Liquid Chromatography–Mass Spectrometry Peptide Feature Detection with Ion Mobility Support. Rapid Commun. Mass Spectrom. 2021, e9045. 10.1002/rcm.9045.

(14) Ivanov, M. V.; Tarasova, I. A.; Levitsky, L. I.; Solovyeva, E. M.; Pridatchenko, M. L.; Lobas, A. A.; Bubis, J. A.; Gorshkov, M. V. MS/MS-Free Protein Identification in Complex Mixtures Using Multiple Enzymes with Complementary Specificity. J. Proteome Res. 2017, 16 (11), 3989–3999. 10.1021/acs.jproteome.7b00365.

(15) Bouwmeester, R.; Gabriels, R.; Hulstaert, N.; Martens, L.; Degroeve, S. DeepLC Can Predict Retention Times for Peptides That Carry As-yet Unseen Modifications. Nat. Methods 2021, 18 (11), 1363–1369. 10.1038/s41592-021-01301-5.

(16) Ivanov, M. V.; Bubis, J. A.; Gorshkov, V.; Abdrakhimov, D. A.; Kjeldsen, F.; Gorshkov, M. V. Boosting MS1-Only Proteomics with Machine Learning Allows 2000 Protein Identifications in Single-Shot Human Proteome Analysis Using 5 Min HPLC Gradient. J. Proteome Res. 2021, 20 (4), 1864–1873. 10.1021/acs.jproteome.0c00863.

(17) Riederer, P.; Nagatsu, T.; Youdim, M. B. H.; Wulf, M.; Dijkstra, J. M.; Sian-Huelsmann, J. Lewy Bodies, Iron, Inflammation and Neuromelanin: Pathological Aspects Underlying Parkinson’s Disease. J. Neural Transm. 2023, 130 (5), 627–646. 10.1007/s00702-023-02630-9.

(18) Ndayisaba, A.; Kaindlstorfer, C.; Wenning, G. K. Iron in Neurodegeneration – Cause or Consequence? Front. Neurosci. 2019, 13. 10.3389/fnins.2019.00180.

(19) Mezzaroba, L.; Alfieri, D. F.; Colado Simão, A. N.; Vissoci Reiche, E. M. The Role of Zinc, Copper, Manganese and Iron in Neurodegenerative Diseases. Neurotoxicology 2019, 74, 230–241. 10.1016/j.neuro.2019.07.007.

(20) Visanji, N. P.; Collingwood, J. F.; Finnegan, M. E.; Tandon, A.; House, E.; Hazrati, L.-N. Iron Deficiency in Parkinsonism: Region-Specific Iron Dysregulation in Parkinson’s Disease and Multiple System Atrophy. J. Park. Dis. 2013, 3 (4), 523–537. 10.3233/JPD-130197.

(21) Dexter, D. T.; Carayon, A.; Javoy-Agid, F.; Agid, Y.; Wells, F. R.; Daniel, S. E.; Lees, A. J.; Jenner, P.; Marsden, C. D. Alterations in the Levels of Iron, Ferritin and Other Trace Metals in Parkinson’s Disease and Other Neurodegenerative Diseases Affecting the Basal Ganglia. Brain J. Neurol. 1991, 114 (Pt 4), 1953–1975. 10.1093/brain/114.4.1953.

(22) Dexter, D. T.; Jenner, P.; Schapira, A. H. V.; Marsden, C. D. Alterations in Levels of Iron, Ferritin, and Other Trace Metals in Neurodegenerative Diseases Affecting the Basal Ganglia. Ann. Neurol. 1992, 32 (S1), S94–S100. 10.1002/ana.410320716.

(23) Shukla, J. J.; Stefanova, N.; Bush, A. I.; McColl, G.; Finkelstein, D. I.; McAllum, E. J. Therapeutic Potential of Iron Modulating Drugs in a Mouse Model of Multiple System Atrophy. Neurobiol. Dis. 2021, 159, 105509. 10.1016/j.nbd.2021.105509.

(24) Juan, C. A.; Pérez de la Lastra, J. M.; Plou, F. J.; Pérez-Lebeña, E. The Chemistry of Reactive Oxygen Species (ROS) Revisited: Outlining Their Role in Biological Macromolecules (DNA, Lipids and Proteins) and Induced Pathologies. Int. J. Mol. Sci. 2021, 22 (9), 4642. 10.3390/ijms22094642.

(25) Levin, J.; Högen, T.; Hillmer, A. S.; Bader, B.; Schmidt, F.; Kamp, F.; Kretzschmar, H. A.; Bötzel, K.; Giese, A. Generation of Ferric Iron Links Oxidative Stress to α-Synuclein Oligomer Formation. J. Park. Dis. 2011, 1 (2), 205–216. 10.3233/JPD-2011-11040.

(26) Ozawa, T.; Paviour, D.; Quinn, N. P.; Josephs, K. A.; Sangha, H.; Kilford, L.; Healy, D. G.; Wood, N. W.; Lees, A. J.; Holton, J. L.; Revesz, T. The Spectrum of Pathological Involvement of the Striatonigral and Olivopontocerebellar Systems in Multiple System Atrophy: Clinicopathological Correlations. Brain 2004, 127 (12), 2657–2671. 10.1093/brain/awh303.

(27) Ishizawa, K.; Komori, T.; Sasaki, S.; Arai, N.; Mizutani, T.; Hirose, T. Microglial Activation Parallels System Degeneration in Multiple System Atrophy. J. Neuropathol. Exp. Neurol. 2004, 63 (1), 43–52. 10.1093/jnen/63.1.43.

(28) Ahmed, Z.; Asi, Y. T.; Sailer, A.; Lees, A. J.; Houlden, H.; Revesz, T.; Holton, J. L. The Neuropathology, Pathophysiology and Genetics of Multiple System Atrophy. Neuropathol. Appl. Neurobiol. 2012, 38 (1), 4–24. 10.1111/j.1365-2990.2011.01234.x.

(29) Gerhard, A.; Banati, R. B.; Goerres, G. B.; Cagnin, A.; Myers, R.; Gunn, R. N.; Turkheimer, F.; Good, C. D.; Mathias, C. J.; Quinn, N.; Schwarz, J.; Brooks, D. J. [11C](R)-PK11195 PET Imaging of Microglial Activation in Multiple System Atrophy. Neurology 2003, 61 (5), 686–689. 10.1212/01.WNL.0000078192.95645.E6.

(30) Song, Y. J. C.; Halliday, G. M.; Holton, J. L.; Lashley, T.; O’Sullivan, S. S.; McCann, H.; Lees, A. J.; Ozawa, T.; Williams, D. R.; Lockhart, P. J.; Revesz, T. R. Degeneration in Different Parkinsonian Syndromes Relates to Astrocyte Type and Astrocyte Protein Expression. J. Neuropathol. Exp. Neurol. 2009, 68 (10), 1073–1083. 10.1097/NEN.0b013e3181b66f1b.

(31) Kübler, D.; Wächter, T.; Cabanel, N.; Su, Z.; Turkheimer, F. E.; Dodel, R.; Brooks, D. J.; Oertel, W. H.; Gerhard, A. Widespread Microglial Activation in Multiple System Atrophy. Mov. Disord. 2019, 34 (4), 564–568. 10.1002/mds.27620.

(32) Hol, E. M.; Pekny, M. Glial Fibrillary Acidic Protein (GFAP) and the Astrocyte Intermediate Filament System in Diseases of the Central Nervous System. Curr. Opin. Cell Biol. 2015, 32, 121–130. 10.1016/j.ceb.2015.02.004.

(33) Piras, I. S.; Bleul, C.; Schrauwen, I.; Talboom, J.; Llaci, L.; De Both, M. D.; Naymik, M. A.; Halliday, G.; Bettencourt, C.; Holton, J. L.; Serrano, G. E.; Sue, L. I.; Beach, T. G.; Stefanova, N.; Huentelman, M. J. Transcriptional Profiling of Multiple System Atrophy Cerebellar Tissue Highlights Differences between the Parkinsonian and Cerebellar Sub-Types of the Disease. Acta Neuropathol. Commun. 2020, 8 (1), 76. 10.1186/s40478-020-00950-5.

(34) Geng, H.; Cao, R.; Cheng, L.; Liu, C. Overexpression of Hepatocyte Cell Adhesion Molecule (hepaCAM) Inhibits the Proliferation, Migration, and Invasion in Colorectal Cancer Cells. Oncol. Res. Featur. Preclin. Clin. Cancer Ther. 2017, 25 (7), 1039–1046. 10.3727/096504016X14813914187138.

(35) Chen, E.; Liu, N.; Zhao, Y.; Tang, M.; Ou, L.; Wu, X.; Luo, C. Panobinostat Reverses HepaCAM Gene Expression and Suppresses Proliferation by Increasing Histone Acetylation in Prostate Cancer. Gene 2022, 808, 145977. 10.1016/j.gene.2021.145977.

(36) Schottlaender, L. V.; Bettencourt, C.; Kiely, A. P.; Chalasani, A.; Neergheen, V.; Holton, J. L.; Hargreaves, I.; Houlden, H. Coenzyme Q10 Levels Are Decreased in the Cerebellum of Multiple-System Atrophy Patients. PLOS ONE 2016, 11 (2), e0149557. 10.1371/journal.pone.0149557.

(37) Foti, S. C.; Hargreaves, I.; Carrington, S.; Kiely, A. P.; Houlden, H.; Holton, J. L. Cerebral Mitochondrial Electron Transport Chain Dysfunction in Multiple System Atrophy and Parkinson’s Disease. Sci. Rep. 2019, 9 (1), 6559. 10.1038/s41598-019-42902-7.

(38) Hsiao, J.-H. T.; Purushothuman, S.; Jensen, P. H.; Halliday, G. M.; Kim, W. S. Reductions in COQ2 Expression Relate to Reduced ATP Levels in Multiple System Atrophy Brain. Front. Neurosci. 2019, 13. 10.3389/fnins.2019.01187.

(39) Mnatsakanyan, N.; Jonas, E. A. The New Role of F1Fo ATP Synthase in Mitochondria-Mediated Neurodegeneration and Neuroprotection. Exp. Neurol. 2020, 332, 113400. 10.1016/j.expneurol.2020.113400.

(40) Yang, Y.; Han, A.; Wang, X.; Yin, X.; Cui, M.; Lin, Z. Lipid Metabolism Regulator Human Hydroxysteroid Dehydrogenase-like 2 (HSDL2) Modulates Cervical Cancer Cell Proliferation and Metastasis. J. Cell. Mol. Med. 2021, 25 (10), 4846–4859. 10.1111/jcmm.16461.

(41) Cheng, Y.; He, X.; Wang, L.; Xu, Y.; Shen, M.; Zhang, W.; Xia, Y.; Zhang, J.; Zhang, M.; Wang, Y.; Hu, J.; Hu, J. [HSDL2 overexpression promotes rectal cancer progression by regulating cancer cell cycle and promoting cell proliferation]. Nan Fang Yi Ke Da Xue Xue Bao 2023, 43 (4), 544–551. 10.12122/j.issn.1673-4254.2023.04.06.

(42) Ruokun, C.; Yake, X.; Fengdong, Y.; Xinting, W.; Laijun, S.; Xianzhi, L. Lentivirus-Mediated Silencing of HSDL2 Suppresses Cell Proliferation in Human Gliomas. Tumour Biol. J. Int. Soc. Oncodevelopmental Biol. Med. 2016, 37 (11), 15065–15077. 10.1007/s13277-016-5402-6.

(43) Ma, S.; Ma, Y.; Qi, F.; Lei, J.; Chen, F.; Sun, W.; Wang, D.; Zhou, S.; Liu, Z.; Lu, Z.; Zhang, D. HSDL2 Knockdown Promotes the Progression of Cholangiocarcinoma by Inhibiting Ferroptosis through the P53/SLC7A11 Axis. World J. Surg. Oncol. 2023, 21 (1), 293. 10.1186/s12957-023-03176-6.

(44) Liu, J.; Zhang, C.; Wang, J.; Hu, W.; Feng, Z. The Regulation of Ferroptosis by Tumor Suppressor P53 and Its Pathway. Int. J. Mol. Sci. 2020, 21 (21), 8387. 10.3390/ijms21218387.

(45) Chen, L.; Mao, L.; Lu, H.; Liu, P. Detecting Ferroptosis and Immune Infiltration Profiles in Multiple System Atrophy Using Postmortem Brain Tissue. Front. Neurosci. 2023, 17. 10.3389/fnins.2023.1269996.

(46) Wakabayashi, K.; Takahashi, H. Cellular Pathology in Multiple System Atrophy. Neuropathology 2006, 26 (4), 338–345. 10.1111/j.1440-1789.2006.00713.x.

(47) Liu, M.; Wang, Z.; Shang, H. Multiple System Atrophy: An Update and Emerging Directions of Biomarkers and Clinical Trials. J. Neurol. 2024. 10.1007/s00415-024-12269-5.

(48) Leńska-Mieciek, M.; Madetko-Alster, N.; Alster, P.; Królicki, L.; Fiszer, U.; Koziorowski, D. Inflammation in Multiple System Atrophy. Front. Immunol. 2023, 14. 10.3389/fimmu.2023.1214677.

(49) Poewe, W.; Stankovic, I.; Halliday, G.; Meissner, W. G.; Wenning, G. K.; Pellecchia, M. T.; Seppi, K.; Palma, J.-A.; Kaufmann, H. Multiple System Atrophy. Nat. Rev. Dis. Primer 2022, 8 (1), 1–21. 10.1038/s41572-022-00382-6.

(50) Jellinger, K. A.; Wenning, G. K. Multiple System Atrophy: Pathogenic Mechanisms and Biomarkers. J. Neural Transm. 2016, 123 (6), 555–572. 10.1007/s00702-016-1545-2.

(51) Kaindlstorfer, C.; Jellinger, K. A.; Eschlböck, S.; Stefanova, N.; Weiss, G.; Wenning, G. K. The Relevance of Iron in the Pathogenesis of Multiple System Atrophy: A Viewpoint. J. Alzheimers Dis. JAD 2018, 61 (4), 1253–1273. 10.3233/JAD-170601.

(52) Byun, Y.; Chen, F.; Chang, R.; Trivedi, M.; Green, K. J.; Cryns, V. L. Caspase Cleavage of Vimentin Disrupts Intermediate Filaments and Promotes Apoptosis. Cell Death Differ. 2001, 8 (5), 443–450. 10.1038/sj.cdd.4400840.

(53) Rawlings, N. D.; Barrett, A. J.; Thomas, P. D.; Huang, X.; Bateman, A.; Finn, R. D. The MEROPS Database of Proteolytic Enzymes, Their Substrates and Inhibitors in 2017 and a Comparison with Peptidases in the PANTHER Database. Nucleic Acids Res. 2018, 46 (D1), D624–D632. 10.1093/nar/gkx1134.

(54) Julien, O.; Zhuang, M.; Wiita, A. P.; O’Donoghue, A. J.; Knudsen, G. M.; Craik, C. S.; Wells, J. A. Quantitative MS-Based Enzymology of Caspases Reveals Distinct Protein Substrate Specificities, Hierarchies, and Cellular Roles. Proc. Natl. Acad. Sci. 2016, 113 (14), E2001–E2010. 10.1073/pnas.1524900113.

(55) Kong, A. T.; Leprevost, F. V.; Avtonomov, D. M.; Mellacheruvu, D.; Nesvizhskii, A. I. MSFragger: Ultrafast and Comprehensive Peptide Identification in Mass Spectrometry–Based Proteomics. Nat. Methods 2017, 14 (5), 513–520. 10.1038/nmeth.4256.

(56) Morishima, N. Changes in Nuclear Morphology during Apoptosis Correlate with Vimentin Cleavage by Different Caspases Located Either Upstream or Downstream of Bcl-2 Action. Genes Cells 1999, 4 (7), 401–414. 10.1046/j.1365-2443.1999.00270.x.

(57) Kragh, C. L.; Lund, L. B.; Febbraro, F.; Hansen, H. D.; Gai, W.-P.; El-Agnaf, O.; Richter-Landsberg, C.; Jensen, P. H. Alpha-Synuclein Aggregation and Ser-129 Phosphorylation-Dependent Cell Death in Oligodendroglial Cells. J. Biol. Chem. 2009, 284 (15), 10211–10222. 10.1074/jbc.M809671200.

(58) Herrera-Vaquero, M.; Heras-Garvin, A.; Krismer, F.; Deleanu, R.; Boesch, S.; Wenning, G. K.; Stefanova, N. Signs of Early Cellular Dysfunction in Multiple System Atrophy. Neuropathol. Appl. Neurobiol. 2021, 47 (2), 268–282. 10.1111/nan.12661.

(59) Kawamoto, Y.; Ayaki, T.; Urushitani, M.; Ito, H.; Takahashi, R. Activated Caspase-9 Immunoreactivity in Glial and Neuronal Cytoplasmic Inclusions in Multiple System Atrophy. Neurosci. Lett. 2016, 628, 207–212. 10.1016/j.neulet.2016.06.036.

(60) Li, F.; Ayaki, T.; Maki, T.; Sawamoto, N.; Takahashi, R. NLRP3 Inflammasome-Related Proteins Are Upregulated in the Putamen of Patients With Multiple System Atrophy. J. Neuropathol. Exp. Neurol. 2018, 77 (11), 1055–1065. 10.1093/jnen/nly090.

(61) Kaji, S.; Maki, T.; Ishimoto, T.; Yamakado, H.; Takahashi, R. Insights into the Pathogenesis of Multiple System Atrophy: Focus on Glial Cytoplasmic Inclusions. Transl. Neurodegener. 2020, 9 (1), 7. 10.1186/s40035-020-0185-5.

